# Inferring the selection window in antimicrobial resistance using deep mutational scanning data and biophysics-based fitness models

**DOI:** 10.1101/189019

**Authors:** Pouria Dasmeh, Anh-Tien Ton, Caroline Quach, Adrian W.R. Serohijos

## Abstract

Mutant-selection window (MSW) hypothesis in antimicrobial resistance implies a range for antimicrobial concentration that promotes selection of single-step resistant mutants. Since the inception and experimental verification, MSW has been at the forefront of strategies to minimize development of antimicrobial resistance (AR). Setting the upper and lower limits of MSW requires an understanding of the dependence of selection coefficient of arising mutations to antimicrobial concentration. In this work, we employed a biophysics-based and experimentally calibrated fitness model to estimate MSW in the case of Ampicillin and Cefotaxime resistance in E.coli TEM-1 beta lactamase. In line with experimental observations, we show that selection is active at very low levels of antimicrobials. Furthermore, we elucidate the dependence of MSW to catalytic efficiency of mutants, fraction of mutants in the population and discuss the role of population genetic parameters such as population size and mutation rate. Altogether, our analysis and formalism provide a predictive model of MSW with direct implications in the design of dosage strategies.

## Introduction

The gain of resistance to antimicrobials is indeed one of the most challenging threats to public health. An effective approach to contain antimicrobial resistance (AMR) is multi-layered, employing knowledge of the mechanisms for gain of resistance, ecological dynamics of bacterial species and eventually treatment regimens and strategies^1^.

Since their first medical use, antimicrobials have been administered at a minimum dosage sufficient to inhibit bacterial growth while keeping hazards minimal. This concentration, termed as minimum inhibitory concentration (MIC)^2,3^, measures susceptibility of bacteria to drugs and is the basis of clinical protocols that compare effectiveness of antimicrobials^4^. The emergence of AMR requires that at least two sub-populations of bacterial species exist, with distinct MICs. The dosage response of bacterial growth to antimicrobials thus defines a concentration window that is above the MIC of wild-type and below the MIC of resistant mutants, i.e., mutant-selection window (MSW) hypothesis^5-7^. The upper limit of this window (i.e., MIC of resistant mutants) is termed as mutant-prevention concentration (MPC)^7^. An effective treatment strategy for AMR sets drug dosage above MPC to inhibit the selection of resistant mutant^6^.

Setting the upper and lower limits of MSW is critically important. Obviously, MPC should be high enough to prevent emergence of AMR, but should be the lowest efficacious concentration to minimize side effects. The lower limit, on the other hand, is the minimum selective concentration (MSC)^8^ below which resistance and efficacy are hardly achieved and can be regarded as a safety level for antimicrobial concentration in natural environments^9^. MSC was traditionally assumed to be the MIC of WT strains, but numerous findings found MSC to be 10- to 100-folds lower and concluded that the selection of resistant mutants is present at sub-MIC concentrations^10^.

Emergence of AMR by drug selection pressure is an example of an evolutionary process whose fate is determined by the competition between mutational supply, selection and immigration^11^. Therefore, the rate of acquired resistance depends on the competition between WT and resistant mutants, which in turn depends on several factors such as the mutation rate, selection regime, population size, etc. In the limit of large population size and low mutation rate, selective advantage of the arising mutation becomes the key factor in determining the evolutionary dynamics. In the simplest definition, the selection coefficient (s) of a mutant is the normalized differential fitness between WT and mutant strains, i.e., F_WT_ and F_mut_:

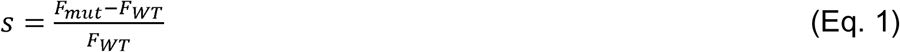

In principle, deleterious, neutral and beneficial nature of mutations is determined by the product of effective population size and selection coefficient, i.e., N_eff_×*s* so that N_eff_× *s* < -1, = 1 and > 1 corresponds to deleterious, neutral and beneficial mutations, respectively.

Over the last decade, considerable efforts have been made to predict protein phenotypes that derive adaptation^12,13^, in particular to drug concentration^14,15^ with the aim of providing an *ab initio* molecular model for AMR. When the gain of resistance is through *de novo* mutations in target proteins^16^, the knowledge of genotype to phenotype mapping is crucial in predicting fitness and eventually the selection regime. In these studies, fitness is proportional to bacterial growth, which depends on the metabolic flux of enzymes targeted by antimicrobials such as *dihydrofolate reductase* (DHFR) in folate pathway^14^ or competitive inhibition in the case of TEM-1 beta lactamase^15,17^.

Given such predictive models of AMR and the resulting fitness landscape from exhaustive mutagenesis experiments, an interesting question is whether these models can be employed to investigate the limits of MSW in AMR. If proved successful, this “bottom-up” approach would result in a predictive model for selective window that will eventually aid in optimizing dosage regimens. This is the major goal of the current work. We will start with a general summary of the predictive model of AR in TEM-1 beta lactamase which relates fitness to the rate of peptidoglycan synthesis and cell wall formation. This rate in turn depends on variety of molecular and cellular factors from permeability of antimicrobials to the catalytic efficiency, k_cat_/K_M_, of both penicillin binding proteins (PBPs) and TEM-1 beta lactamase^17,18^. MSW is then derived from the fitness equation and the landscape for selection coefficients of arising AMR is explored. We then employ this method to investigate the lower and upper theoretical limits of selective window in AMR and comment on dependence of this limit to factors such as enzyme kinetics and the fraction of resistant mutants in the population.

## Results

### Fitness model for TEM-1 beta lactamase

TEM-1 beta lactamase or in general beta lactamases are a family of enzymes that give resistance to beta lactam antibiotics^19-22^. This group of antibiotics which share the common functional group of beta lactam, inhibits formation of the peptidoglycan cross-links in the bacterial cell wall. The specific mode of action of beta lactam antibiotics is the competitive binding with DD-transpeptidases, which are collectively known as penicillin binding proteins (PBP)^23^. PBPs are involved in the final step of peptidoglycan synthesis and thus cell wall biogenesis^24^. Resistance to beta lactam antibiotics, by beta lactamases, is achieved through attacking the beta lactam ring of these antibiotics. Therefore, bacterial growth under the concentration of beta lactam antibiotics is proportional to the activity of PBPs.

Figure 1 depicts the mechanistic model for the function of TEM-1 beta lactamases within the cell. As in Figure 1A, beta lactam antibiotics diffuse through membrane barrier and once in periplasm, they face either hydrolyses by TEM-1 beta lactamases or diffusion towards their target, PBPs. The fitness of a bacterial cell, i.e., growth rate, is modeled by the PBPs activity, v^PBP^ (Figure 1B-C) which is expressed as:

**Figure 1.**
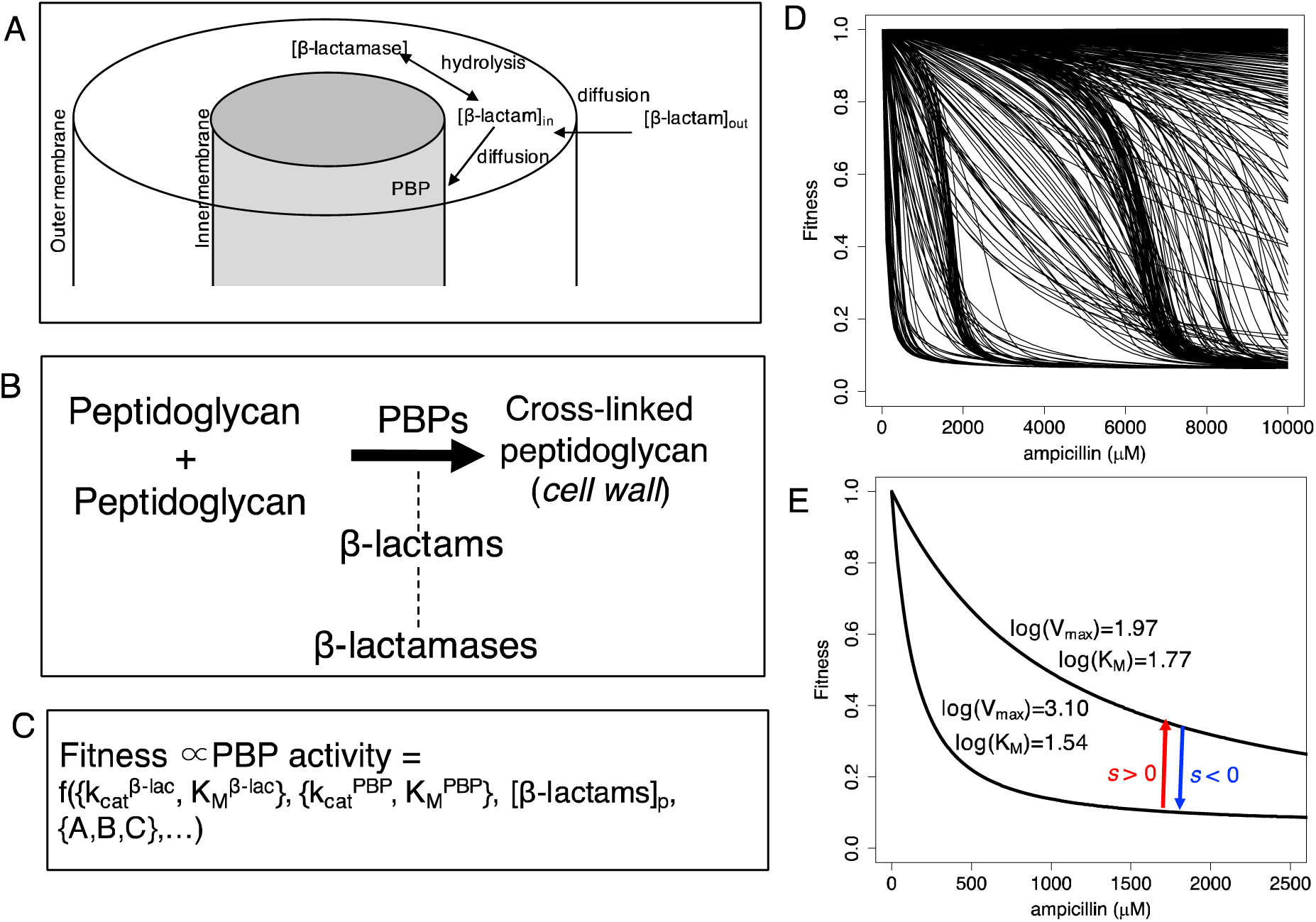
Scheme of the fitness model and the corresponding dosage dependence profile for selection coefficient. A) Beta lactam antimicrobials diffuses through the outer membrane and faces either diffusion to their targets, penicillin binding proteins, PBPs or hydrolysis by beta lactamases. B) Fitness is proportional to the rate of peptidoglycan formation and the activity of penicillin binding proteins. Beta lactams involve in competitive inhibition of PBPs. The resulting fitness function is a function of I) catalytic efficiency of beta lactamases, catalytic efficiency of PBPs, concentration of beta lactams in the periplasm and cellular constants such a permeability of beta lactam antimicrobials. D) Using the functionality of growth rate to different molecular and cellular constants, fitness of different mutants can be expressed as a function of antimicrobial concentration. E) Selection coefficient of an arising mutation can be expressed as a function of antimicrobial concentration depending on the fitness curves of WT and resistant mutants. If mutant is more fit than WT (e.g., has a higher kinetic efficiency-the upper curve), selection coefficient is positive, i.e., red arrow, otherwise negative, i.e., the blue arrow.

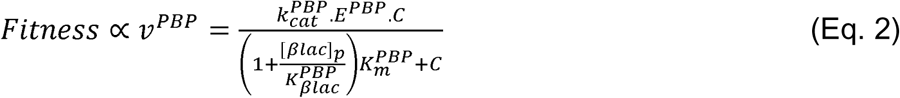

Here, C and [ßlac]_p_ are the periplasmic concentration of peptidyglycan and beta lactam antibiotic, 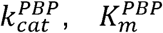, *E*^*PBP*^ and 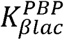 are turnover rate, Michaelis constant, concentration of active PBPs in the periplasm and, dissociation constant of beta lactam antibiotic from PBPs. As assumed in previous works^17,18^, the rate of beta lactam hydrolysis is equal to that of diffusion across periplasm. Hence, one can find the following expression for the concentration of beta lactam antibiotic in the periplasm:

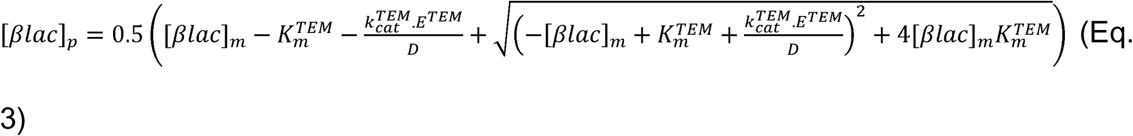

In Eq. 3, [*βlac*]_*m*_ and *D* are the concentration and the permeability constant for beta lactam antibiotic and 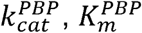, and *E*^*PBP*^ are turnover rate, Michaelis constant and the concentration of active beta lactamases in the periplasm.

Substituting [*βlac*]_*p*_ from Eq. 3 into Eq. 2, selection coefficient (Eq. 1) of an arising mutation can then be calculated as:

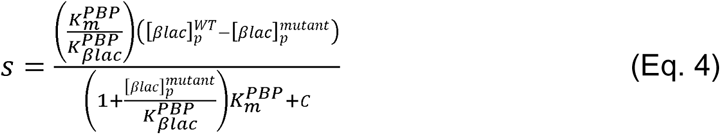

We define the lower and upper limits of MSW, namely MSC and MPC, as the two roots of Eq. 4. We then systematically explore MSC, MPC and MSW as a function of biophysical properties and population genetic parameters. For these analyses, we used three datasets. First, a set of k_cat_ and K_M_ of 4997 TEM-1 mutants against ampicillin that was obtained by fitting the described fitness model to the relative fitness data from a recent mutational scan^17^. Second, two sets of known kinetic data for TEM-1 variants as tabulated in Table S1-2 for reactions with ampicillin and cefotaxime.

We first considered the dependence of selection coefficient to the antimicrobial concentration. Figure 2 shows the average and standard deviation of *s* for all 4997 TEM-1 mutants against Ampicillin. From the figure, mutants become more deleterious i.e., having larger negative selection coefficients, at higher ampicillin concentrations. Figure 2B shows the same plot with a finer range of s = -0.02 to 0.02. The area depicted in gray shows the nearly neutral range (hereafter called nearly-neutral zone) corresponding to |N_eff_×*s*|∼1, where the effective population size, N_eff_, is assumed to be 10^7^. Among 4997 TEM-1 mutants, 32 mutants possess positive selection coefficients at different ranges of antibiotic concentration. As shown in the case of I173H mutant, selection coefficient of beneficial mutations has a quadratic dependence on the antimicrobial concentration and crosses the nearly-neutral zone at a lower and a higher concentration of antibiotic. The lower and higher concentrations correspond to the minimum selective concentration, MSC and, mutant prevention concentration, MPC, respectively (see Methods).

**Figure 2.**
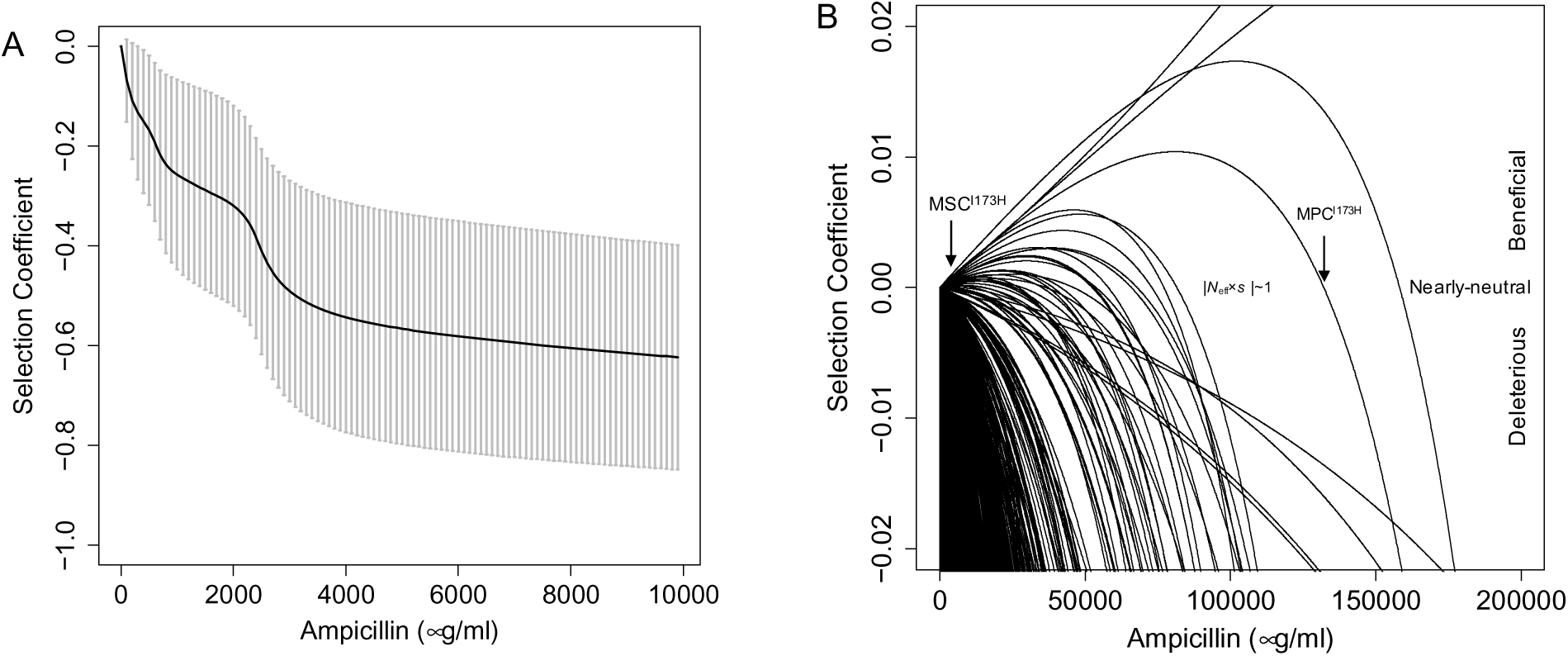
Selection coefficient of TEM-1 mutants versus ampicillin concentration. A) Average (black line) and standard deviation (gray bars) of selection coefficient of 4997 mutants of TEM-1 versus ampicillin concentration. B) Beneficial mutations show a quadratic concave dependence to ampicillin concentrations. The gray area is the zone at which the mutations have nearly-neutral effects on fitness. Above and below this zone, mutations are either beneficial or deleterious. Minimum selective concentration (MSC) and mutant prevention concentration (MPC) are the concentrations at which selection coefficients fall in the nearly-neutral zone.

We calculated MSC and MPC for TEM-1 beneficial mutations against ampicillin in Figure 3A-B. As shown in Figure 3A, MSC is ∼ two-three orders of magnitude lower than MIC of TEM-1 against ampicillin (∼1000-4000 µg/ml ^25-27^). This shows that selection is active at these very low concentrations in line with previous observations for the prevalence of mutant selection at sub-MIC regimes^8,10,28,29^. There is an inverse exponential relationship between MPC and MSC as shown in Figure 3C. Beneficial mutations with higher selective advantages at lower antimicrobial concentrations would indeed require higher antimicrobial concentrations for prevention. The maximum MPC belonged to P27E mutant (∼10^6^ µg/ml). Notably, mutations in residue 173 that are observed in laboratory evolution of TEM-1^30,31^ and in clinical isolates^32^ are among beneficial mutations.

**Figure 3.**
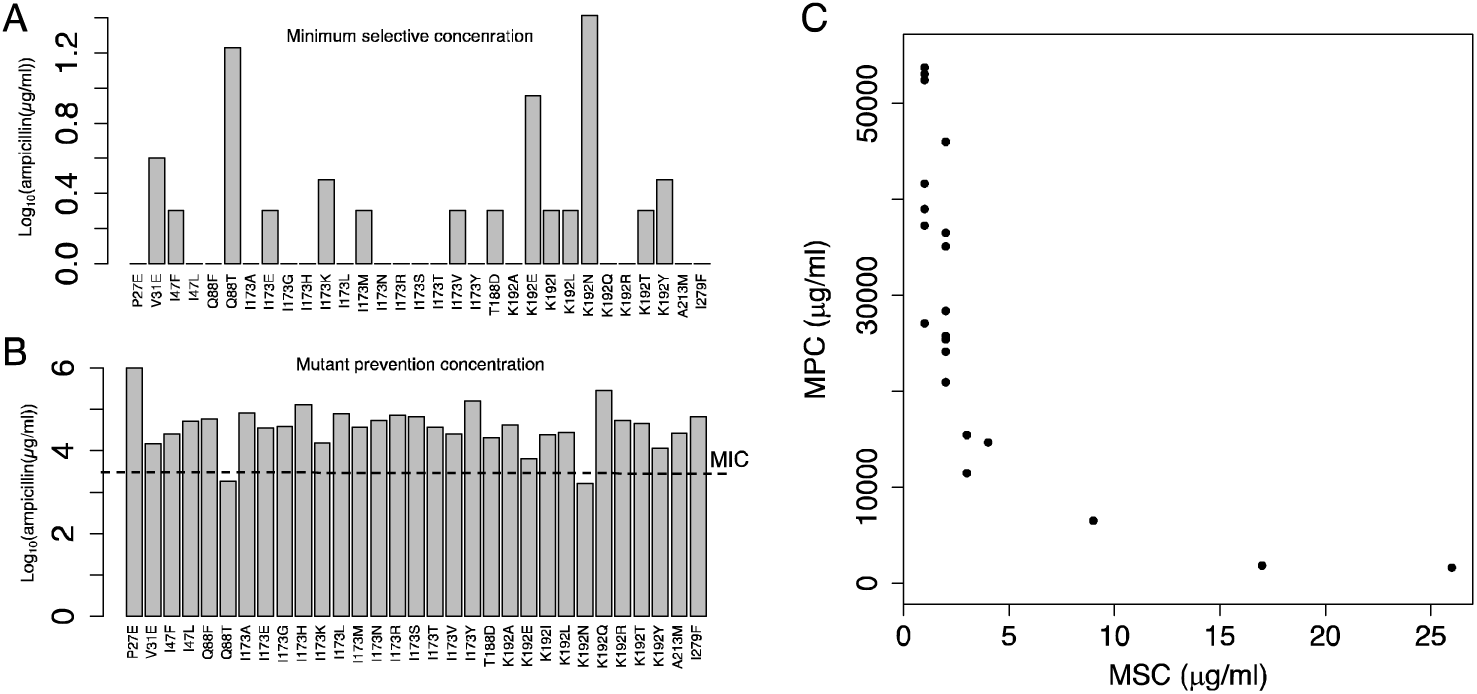
Lower bound, minimum selective concentration, MSC, and upper bound, mutant prevention concentration, MPC of mutant selection window. A) MSC B) MPC and C) MPC versus MSC of beneficial mutants in TEM-1 with ampicillin. The dashed line shows minimum inhibitory concentration, MIC, of WT TEM-1.

Next, we looked at the individual *s* curves for TEM-1 mutants against ampicillin. As shown in Figure 4A, selection coefficients are sigmoidal in antimicrobial concentrations with two interesting features: I) selection coefficients are clustered and non-randomly distributed at different concentrations and II) mutations do not preserve their selective rank at all antimicrobial concentrations. Simply put a high fit mutant to lower concentrations of ampicillin (compared to all other mutants) would not retain the same fitness rank at higher concentrations. Below, we discuss these two features in detail.

**Figure 4.**
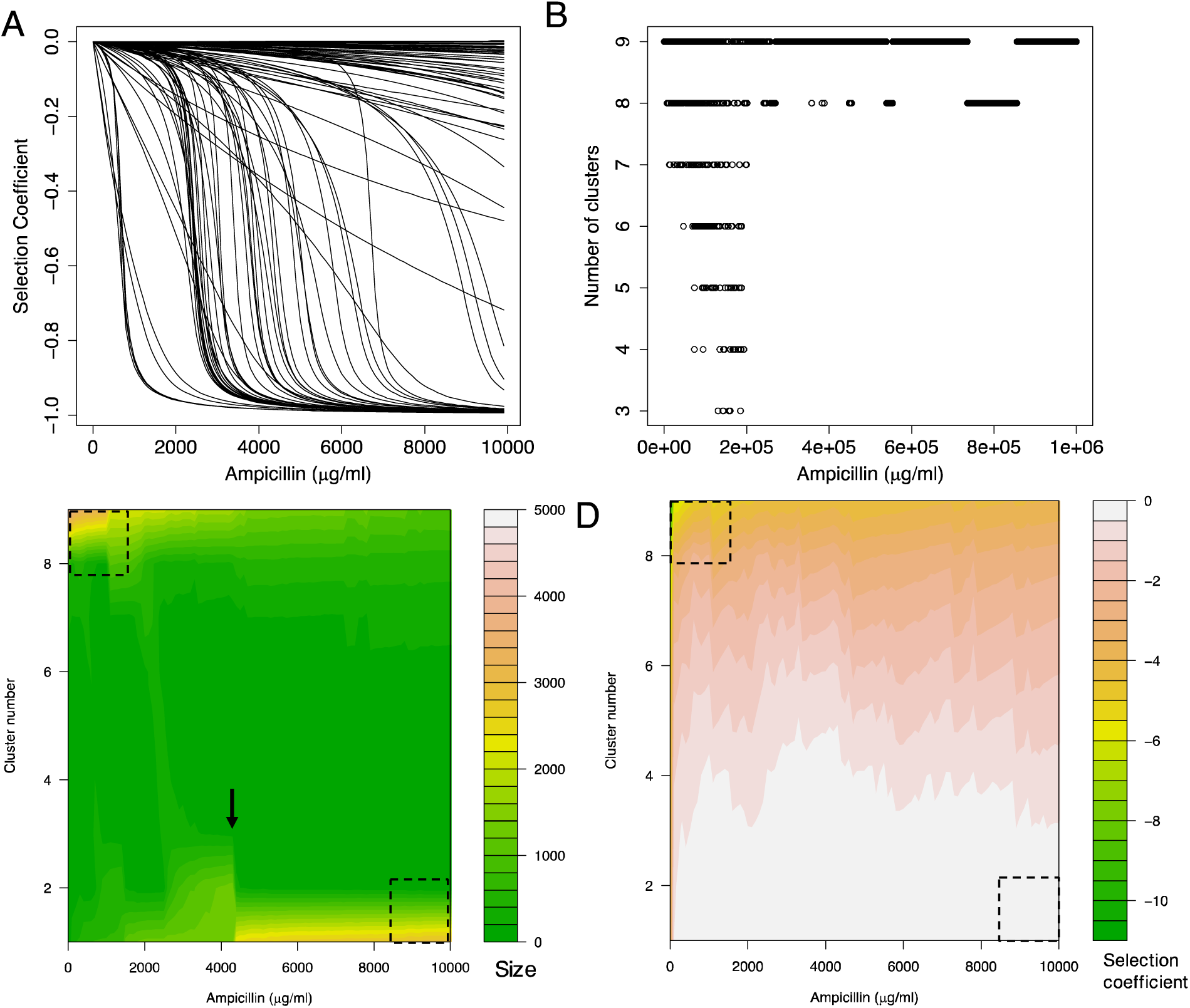
Significant heterogeneity of selection coefficients at different ampicillin concentrations. A) The selection coefficient curves (s-curves) for a set of 100 mutants. B) Clustering of s-curves at different ampicillin concentrations. Up to 9 unique clusters are detected with different sizes. C) Size and D) centroid of each cluster at different antimicrobial concentrations.

The clustering of selection coefficients of several mutations implies that s values are non-uniformly distributed. To see the pattern of clustering, we compared the distribution of distances between selection coefficients with that of an equidistance distribution using Kolmogorov-smirnov test. Selection coefficients at all antimicrobial concentrations were significantly non-equidistant (p-value < 10^-16^). Then, we looked at the optimum number of clusters and the centroids as shown in Figure 4C-D for each ampicillin concentration up to 10000 µg/ml (∼two folds higher than MIC). Several features of these plots are interesting. First, at low antimicrobial concentrations most selection coefficients are within the 8^th^ and 9^th^ clusters being highly deleterious (compare upper left dashed squares in Figures 4C and 4C). This trend is reversed at high concentrations. As shown in bottom-right dashed squares in Figures 4C-D, at high concentrations of antimicrobials, almost half of mutations have neutral or negligible fitness costs. This behavior is in contrast with the average behavior shown in Figure 2A showing a rich heterogeneity in selective advantage/disadvantage of mutant clusters at different antibiotic concentrations. Interestingly, a sudden overpopulation of mutants (∼2/5 of all mutants) in lower more neutral clusters occurs around MIC, shown with a black arrow in Figure 4C.

Another interesting feature of Figure 4A is that s-curves cross each other at different antimicrobial concentrations. This implies that the rank of different mutants is not preserved at all antimicrobial concentrations. To illustrate this observation further, we plotted selection coefficients of two mutants, L152C and G251R, in black and gray in Figure 5A. The corresponding values for V_max_ and K_M_ of both mutants are shown accordingly. From the figure, at low antimicrobial concentrations, the mutant with lower half-saturation limit (L152C) has a higher selective advantage despite having ∼16% less turnover rate. The more efficient G251R mutant gains a higher fitness once antimicrobial concentration reaches the corresponding half-saturation limit.

**Figure 5.**
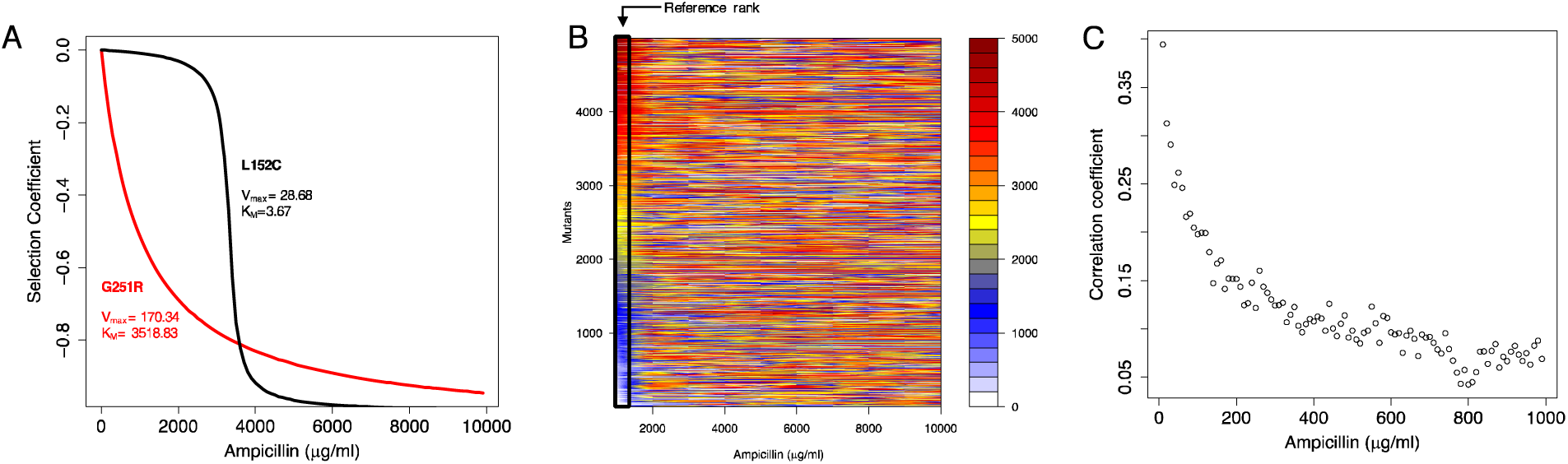
TEM-1 mutants do not preserve their ranks in selective advantage at all concentrations. A) S-curves of two TEM-1 mutants, G251R and L152C, which crosses at antimicrobial concentration of ∼4000 µg/ml. B) Ranks of TEM-1 mutants in selective advantage/disadvantage compared to the rank at 1 µg/ml as the reference set. C) Spearman correlation coefficients between the ranks at each concentration below 1000 µg/ml and the reference at 1 µg/ml.

The lack of rank-preservation at different antibiotic concentrations means that a mutant that confers resistance at one antibiotic concentration would not necessarily do so at a different concentration. This adds another level of impairment for predictability of antimicrobial resistance in addition to stochastic rise and fixation of resistant mutations. To check the rank of mutants at different antimicrobial concentrations systematically, we compared the rank of all TEM-1 mutants at 9 antimicrobial concentrations 1000 to 9000 µg/ml to the ranks at 1 µg/ml. As shown in Figure 5B, the original rank is lost at all higher concentrations. In fact, the correlation between the rank of mutants among all concentrations although significant is very weak (R∼0.01, p-value<10^-3^; see Figure S1-2). We also checked the preservation of ranks at ampicillin concentrations below 1000 µg/ml. As shown in Figure 5C, the maximum correlation is ∼0.35 which decays to ∼0.05 as the antimicrobial concentration increases.

To explore the dependence of MSW to enzyme kinetics, we plotted V_max_ and K_M_ of TEM-1 beneficial mutations. Deleterious and neutral mutations are shown in gray and beneficial mutations in black. From the figure, beneficial mutations have maximized V_max_ and minimized K_M_ leading to maximized V_max_/K_M_ ratio (see the fitted dashed line to all beneficial mutations in Figure 6A). We then plotted MSC and MPC versus V_max_/K_M_ of all beneficial mutants of TEM-1 (Figure 6B-C). From the figure, mutants with a higher catalytic efficiency have lower MSC and higher MPC giving rise to an increased MSW. We thus conclude that V_max_/K_M_ ratio is a strong determinant of MSC and MPC in the case of TEM-1 and ampicillin resistance.

**Figure 6.**
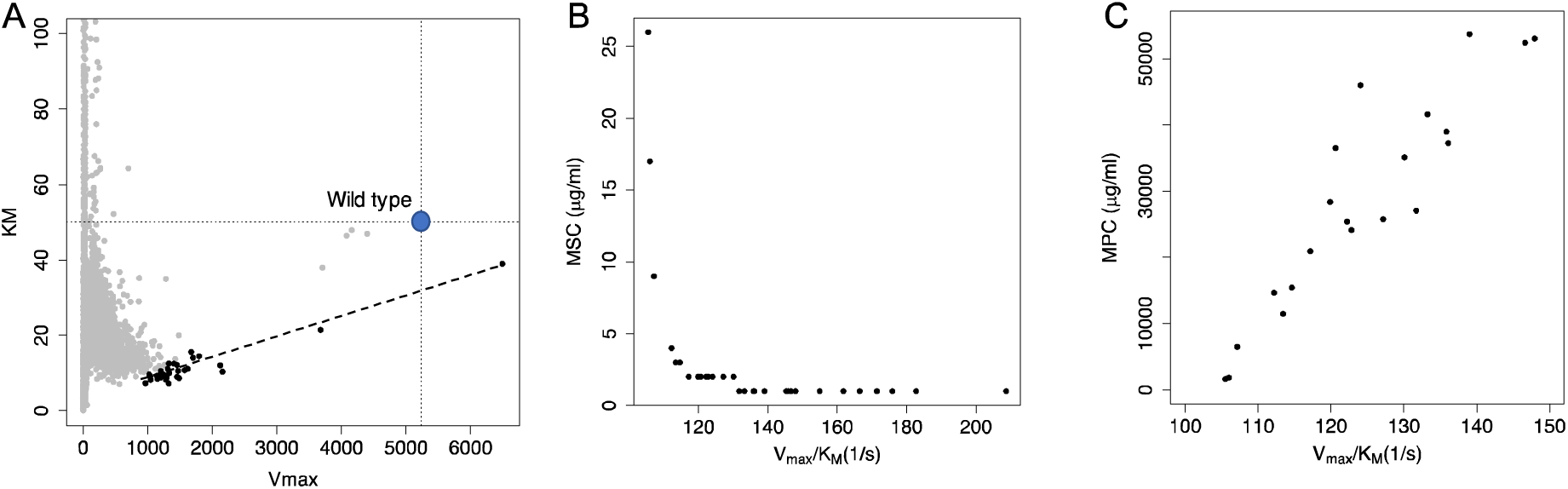
MSC and MPC of TEM-1 beneficial mutations as a function of catalytic efficiency. A) K_M_ versus V_max_ for all (shown in gray) and beneficial (shown in black) TEM-1 mutants. B) MSC and C) MPC of beneficial mutants versus catalytic efficiency.

To what extend the selection regime and the concentration-dependence of selection coefficient discussed in Figure 2 are common to other beta-lactam antimicrobials? To answer this question, we sought to analyze fitness effects of arising mutations in TEM-1 against cefotaxime. Resistance to cefotaxime is widely reported for TEM-1 and the extended-spectrum beta lactamases (ESBLs). Figure S2 shows the selection coefficient curves for TEM-1 mutants and natural variants where kinetic data for reaction with cefotaxime were available (Table S2). Evidently, a major difference with the case of ampicillin is the degree to which mutants are beneficial. While almost all TEM-1 mutants (∼99% in the DMS data set) and all natural variants (Figure S3 and Table S2) were deleterious at all ampicillin concentrations, kinetic data against cefotaxime shows a substantially higher fraction of beneficial mutants (12 out of 16 mutants, Figure S4). From the figure, all ESBLs and TEM-1 mutants share the initial rise in selection coefficient within the few hundred µg/ml of cefotaxime. Note that MIC of cefotaxime is ∼6 µM. The strongly beneficial nature of mutants in TEM-1 against cefotaxime causes MSC to be ∼1 µg/ml for all mutants. MPC is never reached within the concentration range studied here, i.e., up to 10^6^ µg/ml. As shown in Figure S5, the maximum *s* in cefotaxime dataset reaches is almost two orders of magnitude higher than the selection coefficient of the most beneficial TEM-1 mutant against ampicillin.

## Discussion

The knowledge of antimicrobial concentration at which resistance is prevented is crucial for designing dosage strategies. As we showed in this work, and for the specific case of TEM-1 beta lactamase against ampicillin and cefotaxime, a direct determination of MSC and MPC is feasible if the fitness landscape of enzymes conferring antimicrobial resistance, such as TEM-1 beta lactamase is known. As we showed in this work, both MSC and MPC depend on catalytic efficiency of mutants, i.e., the V_max_/K_M_ ratio in the case of ampicillin. However, for strongly beneficial mutations, as observed in the case of cefotaxime, MSC is essentially zero and MPC fall beyond the range of biologically relevant concentrations. One would thus expect that resistance to cefotaxime is much more prevalent than to ampicillin. Therefore, it is essential to estimate not only the fraction of beneficial mutations but also the magnitude of fitness effects, i.e., the full distribution of fitness effects (DFE). We thus propose that DFE obtained from deep mutational scanning combined with fitness models, either biophysics-based as in this work or phenomenological^33^, has a great potential to estimate the spectrum of resistance.

In all calculations in this work, we assumed that both WT and mutants have equal initial finesses. This assumption is biologically unrealistic as resistant mutants are shown to be enriched in the population from very low fractions of the order of 10^-4 29^. To check for this effect, we used the fraction of mutants in the population as a scaling factor for fitness in Equation 2 (see Methods) and plotted selection coefficients versus ampicillin for the case of I173H mutant in Figure 7B. From the figure, MSW is essentially zero for the same beneficial mutation but with negligible fractions in the population. Therefore, figures 7B entails a window of opportunity for containment of AMR if antibiotics are administered at the very onset of emergence of resistance. An exact profiling of both type and fraction of resistant mutants is necessary for this purpose.

**Figure 7.**
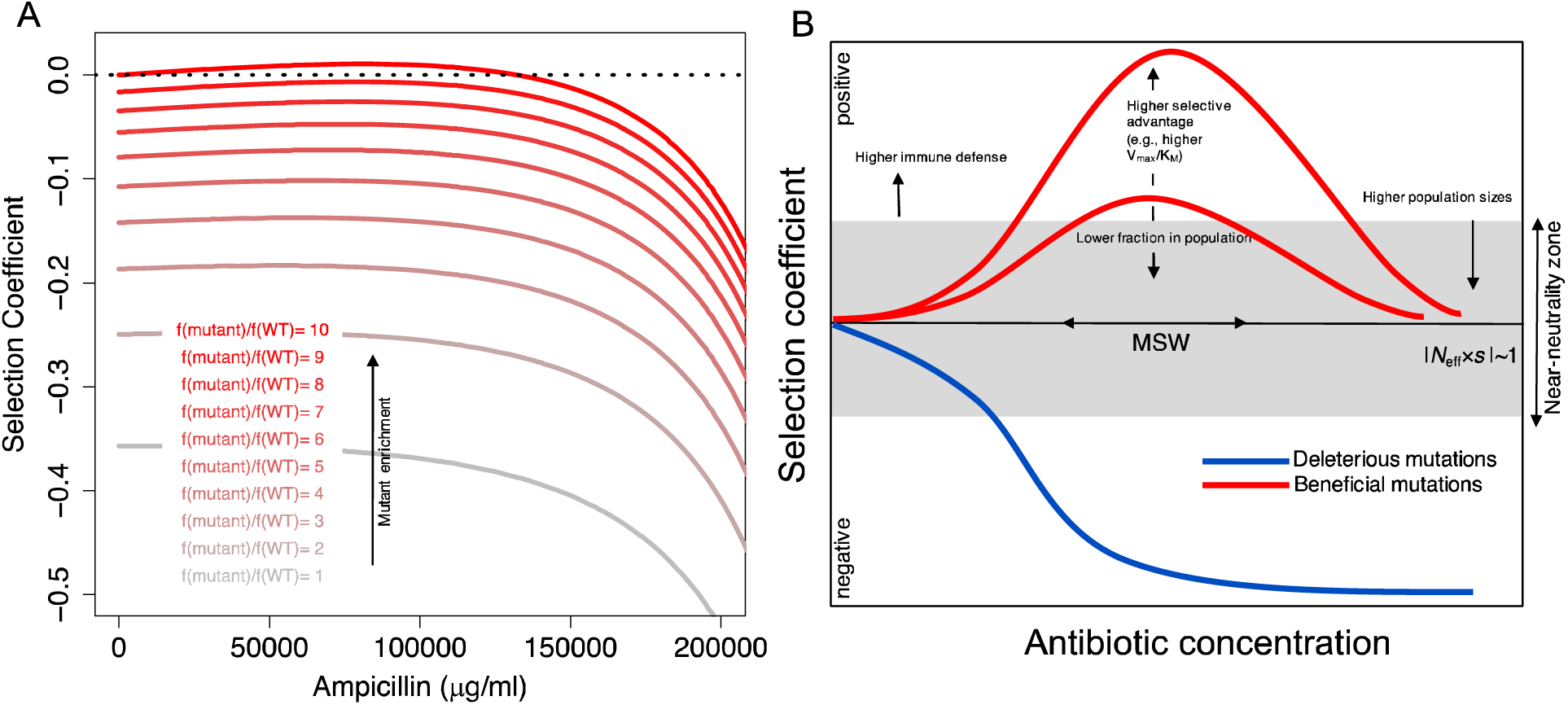
A) Selection coefficient of a beneficial TEM-1 mutant (I173H) versus ampicillin concentration as a function of mutant fraction in the population. B) The schematic influence of confounding factors such as enzymatic efficiency and fraction in the population on MSW.

Two factors, namely bacterial population size and immune response, influence the near-neutrality zone and potentially influence MSW and the fraction of beneficial mutations. First, from classical population genetics, population size scales up fixation of beneficial mutations. Since selection is stringer at higher population sizes, slightly beneficial mutations at a smaller population (e.g., in the regime of N_eff_.s∼1) could be fixated with a higher rate at the larger population. Therefore, higher population sizes decrease the width of near-neutrality zone and gives rise to lower MSC, higher MPC and a wider MSW.

Second, in the absence of immune defense as in laboratory settings, the near-neutrality zone defines the lowest limit of selection. However, a major difference between these settings and the clinical settings as in the patient body is the presence of immune defense. In fact, the first response to bacterial infections is through the immune system. Therefore, one expects that the real selection limit is extended above the near-neutrality zone and to the limit below which infection is contained by the immune defense. This factor narrows down the MSW and thus the number of resistant conferring mutations is further decreased.

We summarize the role of confounding factors on MSW as studied in this work in Figure 7B. First, any factor that increases selective advantage of mutants such as a higher catalytic efficiency widens MSW. We provided an equation for the relationship between MSC and MPC to the V_max_/K_M_ ratio and expect that such relationships exist for other antimicrobials and target enzymes. Second, MSW is substantially narrowed when the fraction of resistant mutations is negligible in the population. Third, lower population sizes or higher immune defense extend the near-neutrality zone and thus narrow down MSW. We anticipate that optimization of MSW with respect to each of these factors would provide a powerful strategy for rational design of dosage strategy in the treatment of antimicrobial resistance.

## Materials and method

### Fitness function

Determination of both upper and lower limits to MSW in our approach requires proper definition of fitness of an organism. Following the original work of Zimmerman and Rosselet^18^ and the following theoretical works^17,34,35^, we assumed that beta lactam antimicrobials passively diffuse through the outer membranes^36,37^. Once in the periplasm, the antimicrobial either get hydrolyzed by beta lactamase or diffuses to the target PBP where competitively inhibits formation of peptidoglycan by interacting with penicillin binding proteins (Figure 1A). The rate equations for beta lactam antibiotics and peptidoglycan synthesis can be written as

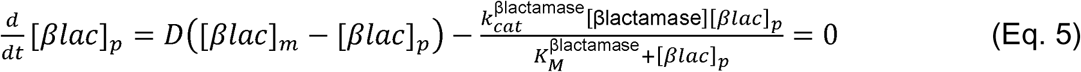

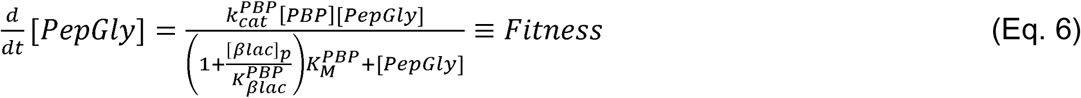

As described earlier, the rate of beta lactam diffusion in the periplasm is assumed to be equal to that of hydrolysis by beta lactamase^17,18^. Using fitness from Eq. 6, the relative fitness 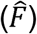 of each mutant to WT TEM-1 is then expressed as:

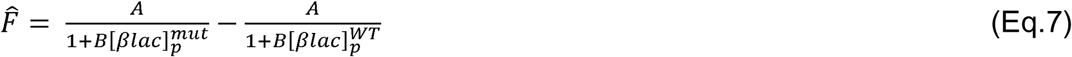

where 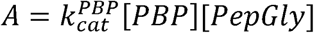 and 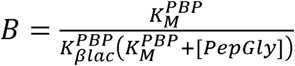. Stiffler et al.^17^ fitted equation 7 to relative fitness obtained from deep mutational scanning data and estimated both A and B constants and K_cat_ and K_M_ of 4997 TEM-1 mutants using a monte carlo simulated annealing approach.

**Table 1.** Parameters used in the fitness function with the corresponding values.

| Parameter | Definition | Value (unit) | References |
| --- | --- | --- | --- |
| $K_{cat}^{PBP}$ | Enzyme turnover rate of PBPs | $\sim 0.05 \text{ (s}^{-1}\text{)}^a$ | 38,39 |
| $K_M^{PBP}$ | Michaelis constant of PBPs | $\sim 83 \text{ mM}$ | 38,39 |
| $E_{PBP}$ | Concentration of folded and active PBPs | $0.12 - 4 \text{ }\mu\text{M}$ | *Calculated*<sup>b</sup> |
| $E_{PG}$ | Periplasmic concentration of peptidoglycan | $\sim 11 \text{ }\mu\text{M}$ | 40 |
| $K_{\beta lac}^{PBP}$ | Dissociation constant of beta-lactam from PBPs | $200 - 1600 \text{ }\mu\text{M}^{-1}$ | 41 |
| $k_{cat}^{TEM-1}$ | Enzyme turnover rate of TEM-1 | $\sim 1 - 5000 \text{ (1/s)}$ | 25,26,42-50 |
| $K_M^{TEM-1}$ | Michaelis constant of TEM-1 | $\sim 4 - 3500 \text{ (1/s)}$ | 25,26,42-50 |
| $E_{TEM-1}$ | Concentration of folded and active TEM-1 | $\sim 3 - 4 \text{ }\mu\text{M}$ | 17 |
| $D$ | permeability parameter for beta-lactam | $\sim 0.002 - 0.005$ | 17 |

a: For values in ranges, the average value is used in the model. b: See supplementary information for details.

